# Transport-driven spatial patterning of glucosinolates structures root microbiome assembly

**DOI:** 10.64898/2026.02.23.707459

**Authors:** Andra-Octavia Roman, Meike Burow, Lioba Rüger, Tonni Grube Andersen

## Abstract

Plant roots actively assemble distinct microbial communities, yet how host chemical traits are organized to structure them remains poorly understood. Glucosinolates are hallmark defense metabolites of Brassicaceae, but their axial distribution in roots and ecological relevance belowground remain largely unknown. Here, we combine spatial metabolite profiling and microbiome analysis in *Arabidopsis thaliana* and the oilseed crop *Camelina sativa* using mutants lacking the glucosinolate transporters GTR1 and GTR2. We find that both species exhibit a conserved, transporter-dependent enrichment of long-chained aliphatic glucosinolates at the root tip, revealing active axial organization of chemical defenses in roots. Using 16S rRNA amplicon-based sequencing, we show that plant species identity is the primary determinant of bacterial community composition. However, disruption of axial glucosinolate distribution significantly alters spatial patterns of microbiome assembly along the root in a species-dependent manner. In *Arabidopsis*, this assembly effect is most pronounced in the rhizosphere, whereas in *Camelina,* root-associated communities were also affected. Together, our findings demonstrate that glucosinolate transport establishes chemical landscapes along the root axis, thereby shaping spatial patterns of microbiome assembly. This identifies spatially structured specialized metabolite allocation as an important mechanism by which plants shape their belowground microbial environment.

## Introduction

Plants have evolved sophisticated mechanisms to interact with other organisms, particularly in the microbe-rich soil surrounding their roots. To persist in this environment, they mediate chemical interactions with microbes [1]. This enables plants to limit harmful colonizers and select for microbes that contribute to root health and ecological stability [2]. A key component of this strategy is active recruitment and structuring of a beneficial root-associated community or microbiota [3–5]. A central aspect of plant-microbe interactions is the production of specialized metabolites, which can serve as defensive compounds that actively kill or inhibit colonization by specific microbes [6–8].

One intriguing aspect of root-associated plant–microbe associations is the organization across the root system. This occurs across radially distinct rhizocompartments: the rhizosphere (the soil immediately surrounding the root), the rhizoplane (the root surface), and the endosphere (the internal tissues) [9]. While the influence of specialized metabolites across these compartments has been clearly demonstrated [10–12], more recent work emphasizes the importance of differences along the root axis [13, 14]. In maize, for instance, root border cells and mucilage secreted from the root tip shape the composition of *Cercozoa, Bacteria*, and *Archaea* in the vicinity of the meristematic region [15, 16]. In *Arabidopsis thaliana* (hereafter *Arabidopsis)*, perturbations in expression of transporters suggest shifts in bacterial community composition specific to distinct longitudinal root sections [17]. Thus, active, precise allocation of metabolites along the root axis may play a role in maintaining niche-specific root–microbe interactions. This introduces an additional dimension for host control of microbial assembly. Yet, for specialized metabolites, it remains unresolved whether such spatial organization reflects passive consequences of biosynthetic activity or constitutes an actively maintained transport-dependent mechanism.

Among specialized metabolites, glucosinolates (GSLs) stand out as arguably the best-characterized group with demonstrated roles in plant defenses [18]. GSLs are sulfur-and nitrogen-containing compounds produced by a limited number of angiosperm families within the order *Brassicales,* which also includes *Arabidopsis* [19]. Over 130 distinct GSLs have been identified [19]. These are grouped into three classes, based on their amino acid precursors: phenylalanine or tyrosine for benzenic GSLs, tryptophan for indolic GSLs, and methionine, alanine, leucine, or isoleucine for aliphatic GSLs, respectively [20–22]. Aliphatic GSLs synthesized from chain-elongated methionine show species-specific variation in side-chain length. Short-chained (SC) aliphatic GSLs (3 to 5 carbons) constitute the majority in *Arabidopsis (*Col-0), with relatively few long-chained (LC) forms (6–8 carbons). Longer-chained GSLs (9 or more carbons) are characteristic of other species, such as the emerging oil crop *Camelina sativa* (hereafter *Camelina*) [20, 23, 24]. These three classes not only differ in biosynthetic origin and abundance, but also in spatial localization and accumulation patterns, depending on plant species and developmental stage [25]. This structural and distribution diversity makes GSLs ideal candidates for studying how specialized metabolites spatially modulate plant–microbe interactions locally.

In leaves of *Arabidopsis*, GSL concentrations are highest in younger tissues or midvein lamina [26–28]. This pattern likely serves to deter herbivores from feeding on younger, more nutritious tissues [26, 28]. Aliphatic GSLs also influence microbial communities in the phyllosphere, contributing to species- and tissue-specific bacterial recruitment [29]. In roots, changes in the presence or absence of specific GSLs, such as indolic GSL, have been shown to affect bacterial community composition [30]. The individual compounds vary between cultivars, as demonstrated by comparisons between laboratory accessions and wild relatives [29]. However, the spatial distribution of GSLs and their specific contribution in distinct root regions remain unresolved.

Genetic resources in Arabidopsis mutants enable dissection of GSL biosynthesis and transport, but only a few are suitable for studying their distribution [25, 31–36]. Key players in this function are GLUCOSINOLATE TRANSPORTER 1 and 2 (GTR1 and GTR2), which mediate GSL allocation between tissues [35, 37], including leaves and roots [32, 38]. In *Arabidopsis*, a *gtr1gtr2* double mutant fails to retain GSLs from the root xylem, which leads to accumulation of root-synthesized LC aliphatic GSLs at the leaf margins instead of in their designated root cells [35]. This mutant has been instrumental in providing insights into mechanisms that underlie the role of GSL transport dynamics in herbivore feeding patterns [26], yet the implications of altered chemical defense organization for root-associated biota remain unknown.

In this study, we investigated the role of GSL distribution in shaping the root and rhizosphere microbiome, focusing on how disruption of their accumulation patterns affects bacterial communities in a natural soil environment. Using GSL transport-disturbed mutants in *Arabidopsis* and *Camelina,* we performed a comparative analysis of both metabolite distribution and bacterial community assembly along the longitudinal root axis. Our findings reveal that bacterial communities differ not only between plant species but also in association with GTR-dependent enrichment of LC aliphatic GSLs along the root axis. This highlights the importance of local allocation of specialized metabolites in underground environments as well as the relevance of root zonation, when evaluating plant–microbe interactions in roots.

## RESULTS

### GTR1 and GTR2 control glucosinolate accumulation in soil-grown *Arabidopsis* and Camelina roots

In *Arabidopsis*, the GSL transporters GTR1 and GTR2 are key determinants of root GSL content under artificial growth conditions [32, 38]. As a *gtr1gtr2* double mutant was recently generated in *Camelina sativa* [39], we set out to test whether the function of these transporters in mediating shoot–root GSL partitioning is conserved across species in soil. We cultivated *Arabidopsis* and *Camelina* wild type and *gtr1gtr2* mutants in CD-cover-based rhizotrons, which enable gentle extraction of intact roots from soil-grown plants [40]. When grown on Cologne agricultural soil (CAS) [41, 42], *gtr1gtr2* mutants exhibited elevated levels of aliphatic GSLs in the shoots of both species compared to wild type; however, the difference reached statistical significance only in Arabidopsis (Figure 1). Consistent with previous findings on agar [35], this difference was predominantly due to an increase in LC aliphatic GSL content (Supp. Figure 1). In roots of both species, *gtr1gtr2* mutants had strongly reduced GSL levels (Figure 1), indicating that their diminished ability to retain GSLs in the root tissue also occurs in *Camelina*. While *Arabidopsis* mutants also displayed changed levels of indolic GSLs in these conditions, *Camelina* does not produce this class of GSL [23, 24] (Supp. Figure 1).

**Figure 1.**
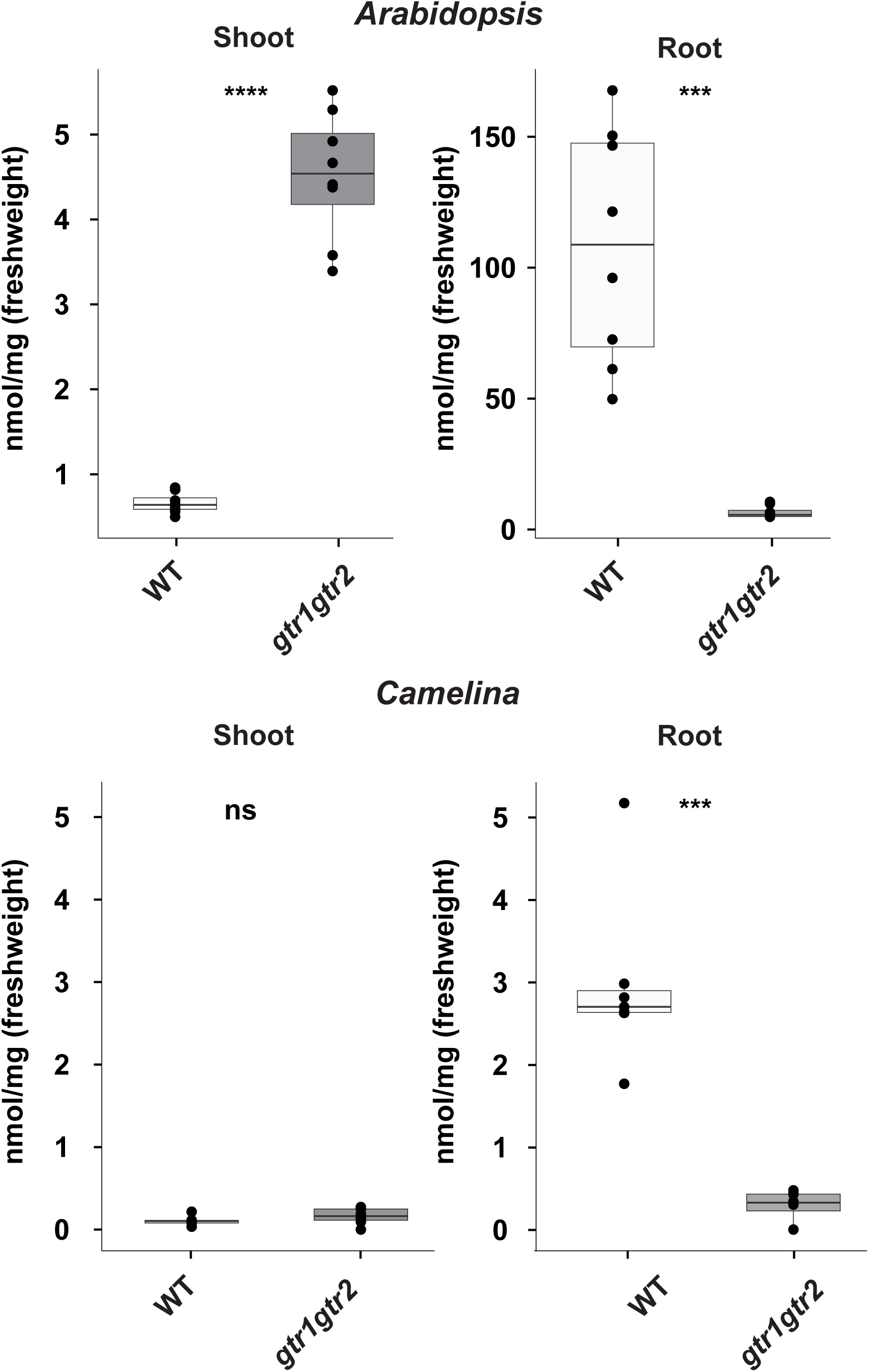
Glucosinolate content in plants grown in Cologne agricultural soil. Concentration of long-chained (LC) aliphatic glucosinolates in tissues of wild type (WT) and *gtr1gtr2* mutants from *Arabidopsis thaliana* (*Arabidopsis*) and *Camelina sativa* (*Camelina*) grown in Cologne agricultural soil. For boxplots, the center line indicates median, dots represent data points, the box limits represent the upper and lower quartiles, and whiskers indicate maximum and minimum values. *** indicates P<0.01 in a Students t-test, n=8; n.s., not significant

### Whole-root analysis reveals species-specific microbiome assembly without detectable genotype effects

Next, to test whether the belowground bacterial community composition is affected in *gtr1gtr2* mutant plants, we performed 16S rRNA amplicon sequencing on root (endosphere and rhizoplane) and rhizosphere fractions of plants grown in CAS. Samples from *Camelina* and *Arabidopsis* were processed and sequenced separately. After filtering ASVs with fewer than 100 reads, we retained 4,709 ASVs representing 7,535,983 sequences in *Arabidopsis* and 3,860 ASVs comprising 5,319,005 sequences in *Camelina*. Rarefaction analyses confirmed sufficient sequencing depth (Supp. Figure 2). Overall, *Arabidopsis* harbored greater ASV richness than *Camelina*. Neither alpha nor beta diversity differed significantly between wild type and *gtr1gtr2* mutants in either species (Figure 2A; Supp. Figure 3; Supp. Table 1). In both species, the alpha diversity was consistently higher in the rhizosphere than in the root compartment, indicating selective colonization of the root (Supp. Figure 3). A principal coordinates analysis (PCoA) and PERMANOVA based on Bray–Curtis dissimilarities further supported strong compartmentalization of bacterial communities, with clear separation of root and rhizosphere samples in *Arabidopsis* (R² = 0.50, P = 0.001) and *Camelina* (R² = 0.38, P = 0.001) (Figure 2A). Relative abundance profiles revealed species-associated compositional differences on bacterial phylum level (Figure 2B; Supp. Table 2). Differences in the relative abundance of bacterial phyla between species were generally stronger in the rhizosphere compared to the root compartment, likely reflecting higher within-group variation in the root compartment. *Arabidopsis*-associated bacterial communities were dominated by *Proteobacteria*, whereas *Camelina* harbored a comparatively higher proportion of *Actinobacteriota* (Figure 2B; Supp. Table 2). Taken together, root compartment and plant species identity explain most variation at the whole-root level in the root-associated bacterial community composition in our tested soil system. This apparent absence of genotype effects at whole-root resolution highlights the potential for spatially confined chemical gradients to influence microbial assembly in ways that are masked when roots are analyzed as homogeneous units.

**Figure 2.**
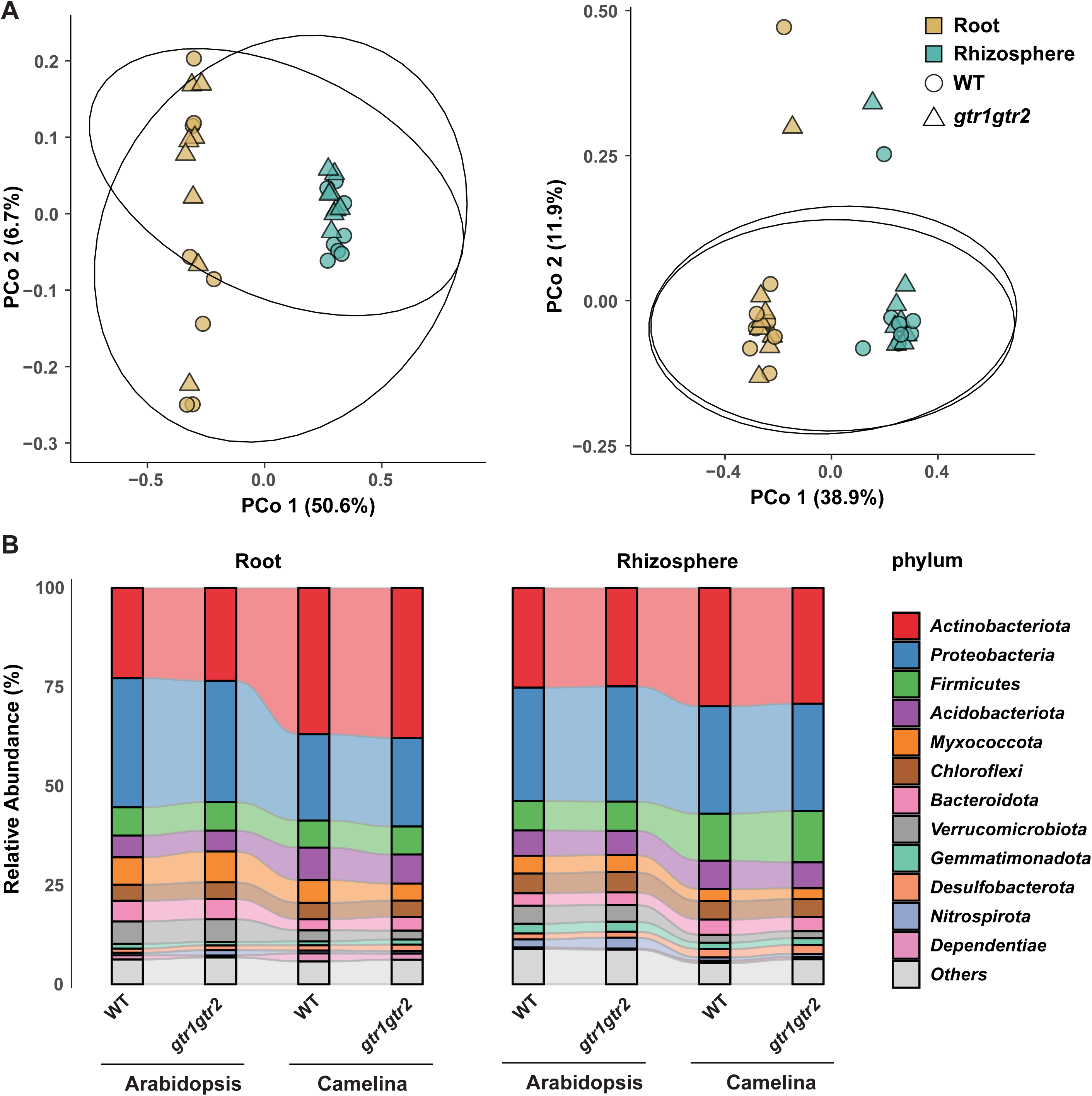
Bacterial community structure in roots of *Arabidopsis* and *Camelina*. A. PCoA of Bray–Curtis dissimilarities of root associated communities in *Arabidopsis* (left panel) and *Camelina* (right panel) showing variation by rhizo-compartment and genotype (n=8). Ellipses represent 95% confidence intervals around centroids of genotype groups. B. Relative abundance of the 12 most abundant bacterial phyla in root (left) and rhizosphere (right) communities (n=8).

**Figure 3.**
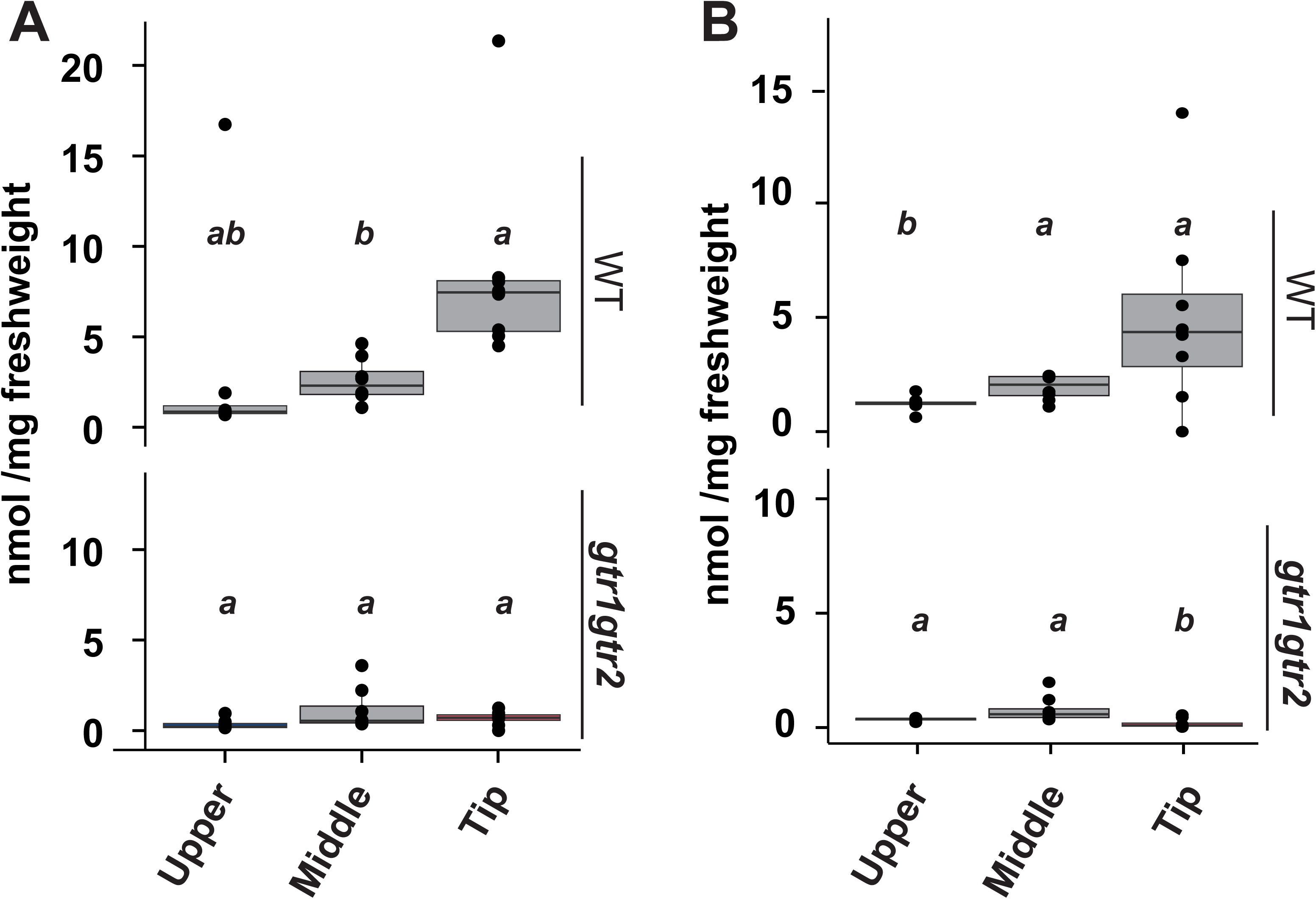
GSL distribution along the root sections in *Arabidopsis* and *Camelina*. LC aliphatic GSL concentrations in upper, middle and tip of the root in wild type and *gtr1gtr2* mutants of A. *Arabidopsis* and B. *Camelina* plants (n=8). Statistical analysis was performed using one-way ANOVA followed by Tukey’s post hoc test (α = 0.05).

### LC aliphatic GSLs accumulate in a GTR-dependent manner in young root regions of Arabidopsis and Camelina

Because GSLs are spatially patterned in leaves [28, 43, 44], we hypothesized that their distribution along the root axis may likewise be actively organized rather than homogeneous. To test this, we quantified GSL content in three distinct root zones: the root tip, middle, and upper segments, which represent developmental transitions from the meristematic region through a zone with lateral root initiation to mature tissues with periderm formation [45]. These regions differ locally in surface characteristics and barrier composition [46] and are therefore candidates to show differences in GSL accumulation.

Indeed, in wild type *Arabidopsis*, LC aliphatic GSLs were most abundant in the root tip (Figure 3A), whereas indolic GSLs accumulated predominantly in upper root segments (Supp. Figure 4). SC aliphatic GSLs were only detected in trace amounts and restricted to the upper region (Supp. Figure 4). In the *gtr1gtr2* mutant, LC aliphatic GSL levels were strongly reduced in the tip region (Figure 3A), while indolic GSL accumulation remained similar to wild type (Supp. Figure 4). Likewise, in *Camelina*, LC GSLs showed the highest accumulation near the root tip in a GTR1/GTR2-dependent manner (Figure 3B). Thus, in soil, GTR-mediated transport governs the enrichment of LC aliphatic GSLs in young root regions, a conserved feature of both Arabidopsis and Camelina, despite their divergent GSL profiles. This suggests that axial allocation represents an evolutionarily maintained feature rather than a species-specific idiosyncrasy.

**Figure 4.**
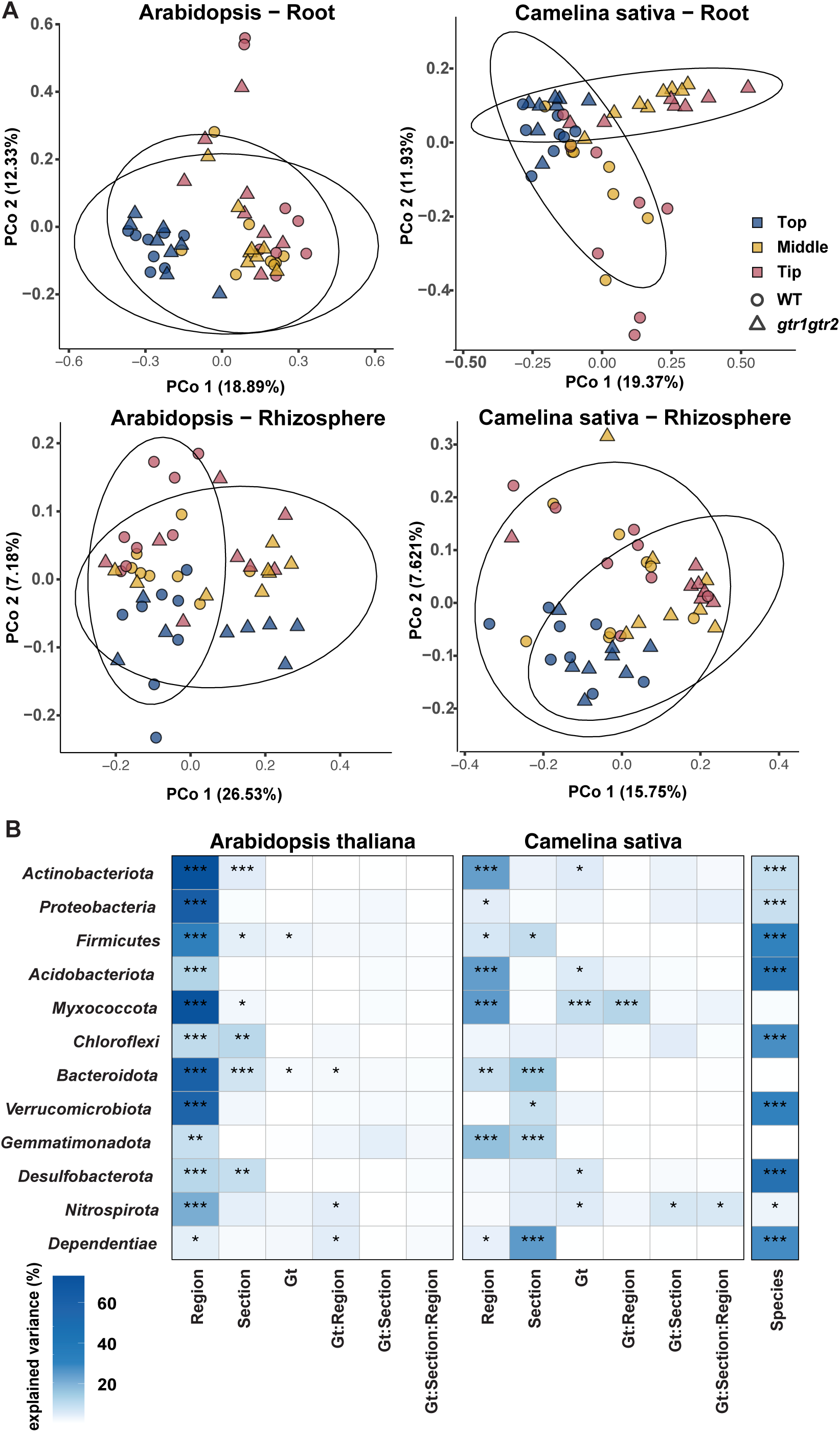
Variation in bacterial community structure in roots of *Arabidopsis* and *Camelina* across rhizo-compartment, root sections, and genotype. A. PCoA of Bray–Curtis dissimilarities of root and rhizosphere associated communities of *Arabidopsis* and *Camelina*, showing variation by root section and genotype (n=8). Ellipses represent 95% confidence intervals around genotype centroids. B. Heatmap showing variance explained and significance (ANOVA) of plant species, rhizo-compartment, root section and genotype (Gt) for the relative abundance of the 12 most abundant bacterial phyla (n=8). Significance is indicated by asterisks: P<0.05 (*), P<0.01 (**), P<0.001 (***).

### Root bacterial communities exhibit GSL-dependent spatial assembly

Motivated by this distinction in GSL distribution, we next asked whether the spatial patterning of GSLs in roots influences microbial community composition along the root. Using *gtr1gtr2* mutants, we profiled bacterial communities from corresponding root zones (tip, middle, upper) and their adjacent rhizospheres of both species grown in CAS. Also in this experiment, the alpha diversity differed between plant species and compartments but not between genotypes (Supp. Figure 5, Supp. Table 1). Beta diversity analyses revealed a more nuanced view of this spatial structuring (Figure 4). In *Arabidopsis*, bacterial communities within the root compartment showed significant differentiation by root section (R² = 0.18, P = 0.001), but not by genotype, suggesting that GSL distribution has limited influence on endophytic colonization. In contrast, in the *Arabidopsis* rhizosphere, both root section (R² = 0.07, P = 0.018) and genotype (R² = 0.09, P = 0.001) significantly explained variance in community composition. This pattern was reflected by partially distinct clusters in the PCoA (Figure 4A), consistent with spatially distributed GSLs contributing to rhizosphere community differentiation. In *Camelina*, both root section and genotype significantly influenced beta diversity across compartments, with significant effects observed in roots (section: R² = 0.14, P = 0.001; genotype: R² = 0.08, P = 0.001) and rhizosphere (section: R² = 0.10, P = 0.001; genotype: R² = 0.05, P = 0.003) (Figure 4A). Pairwise PERMANOVA identified genotype-dependent differences specifically in the tip and middle sections of both species (Supp. Table 1). Together, these results support a model where microbial community assembly along the root axis is spatially structured and partially dependent on axial GSL distribution. While these patterns do not demonstrate direct causality, their spatial concordance with GTR-dependent LC GSL enrichment supports a model in which transporter-mediated allocation constrains microbial assembly along the root axis. However, the influence of GTR-mediated processes differs between hosts, as it impacts the rhizosphere microbiome in *Arabidopsis*, but extends into root- and rhizoplane-associated communities in *Camelina*.

### Distinct bacterial phyla underlie spatial GSL-dependent responses

Comparison of the relative abundance of the 12 most dominant bacterial phyla revealed substantial variation across plant species, root sections, and rhizo-compartments (Figure 4B, Supp. Figure 6; Supp. Table 2). When assessing genotype effects, significant differences were observed only in *Camelina* roots. In particular, the relative abundance of *Myxococcota* in the middle and tip sections and *Nitrospirota* in the middle section differed significantly between wild type and *gtr1gtr2* mutant plants (Supp. Figure 6; Supp. Table 2). To further disentangle the contributions of plant species, spatial position, and genotype, we applied an ANOVA-based model to evaluate their effects on bacterial phyla relative abundances. Plant species, compartment, and root section emerged as the strongest determinants of bacterial phyla composition (Figure 4B). The *gtr1gtr2* mutations had only a minor effect on the most abundant phyla in *Arabidopsis*, but significantly affected *Firmicutes* and *Bacteroidota*, and a genotype–compartment interaction was detected for *Nitrospirota* and *Dependentiae*. In contrast, in *Camelina*, the mutations influenced the relative abundance of several phyla, including *Actinobacteriota*, *Acidobacteriota*, *Desulfobacterota*, *Nitrospirota*, and most strongly *Myxococcota*. Higher-order interactions, specifically those between genotype and compartment, significantly affected the relative abundance of *Myxococcota*. In conclusion, the spatial distribution of GSLs mediated by GTRs is important for shaping the relative abundance of specific bacterial phyla in a species- and compartment-dependent manner, with particularly pronounced effects in *Camelina*.

## Discussion

Root-associated microbiomes are shaped by a complex interplay of host traits, metabolites, and structure [47]. Our data demonstrate a spatial concordance between transporter-dependent LC aliphatic GSL enrichment and compartment-specific shifts in bacterial community composition; however, they do not resolve whether these effects arise from altered exudation, intracellular filtering, or microbial sensitivity to distinct GSL chemistries. Several studies have established that exuded chemicals shape the composition of root-associated microbial communities (*46*–*50*). Thus, the exudation profiles or activation strategies of GSLs may differ between Arabidopsis and Camelina. In contrast, *Camelina*, with its thicker cortex and distinct developmental anatomy, may provide a more selective internal environment, whereas Arabidopsis may exert comparatively stronger control through metabolite release at the rhizosphere interface [45, 53]. These anatomical differences are consistent with our observations but remain speculative, as direct measurements of exudation dynamics or barrier permeability were not performed in this study.

Another intriguing difference between our employed species is that *Arabidopsis* accumulates LC aliphatic GSLs with up to 8 carbon atoms in the side chain ([27], Supp. Figure 1), whereas *Camelina* predominantly accumulates those with 9 to 11 carbons [23, 24]. Thus, certain bacteria may be sensitive to GSLs with longer side chains, which could explain some of the species-specific accumulation patterns observed (Figure 2) [54]. Moreover, *Arabidopsis* produces both LC aliphatic and indolic GSLs, which accumulated in distinct zones of the root (Figure 3, Supp. Figure 4) [55].

Taken together, we conclude that GSL distribution along the root axis is not a passive consequence of biosynthesis, but rather reflects actively maintained, transporter-dependent spatial organization, that likely constrains microbial community assembly. Whereas in Arabidopsis, this may primarily influence microbial assembly at the rhizosphere-root interface, in Camelina it appears to combine deeper internal filtering with longer-chain GSL chemistry. From an ecological perspective, such transporter-mediated allocation of specialized metabolites represents a flexible mechanism by which plants tune microbial filtering across niches without altering biosynthetic capacity.

This interpretation is supported by our finding that spatial differentiation in community composition was greater within roots compared to the rhizosphere, consistent with previous work showing that the endosphere is subject to stronger fine-scale heterogeneity and selective filtering [1]. In contrast, the rhizosphere, where microbial diversity is higher and DNA from dead material can persist, dilutes fine-scale host effects [58]. Against this background, it is particularly striking that in *Arabidopsis* the *gtr1gtr2* mutation altered communities primarily outside roots, suggesting a primary role of LC aliphatic GSL release dynamics in shaping rhizosphere assemblages. The soil surrounding roots includes not only neighboring plant species, but also a wide range of prokaryotes and eukaryotes, such as bacteria, protists, fungi, and invertebrates [59, 60]. While our focus here is on bacterial communities, other groups are likely subject to similar metabolite-mediated filtering. Particular attention has been paid to the impact of metabolites on bacterial communities, both in the rhizosphere and within plant tissues, where both endophytic and rhizosphere-dwelling bacteria contribute significantly to plant health and development [6, 48, 61].

One additional important caveat of our study lies in the difference in plant age between the two species compared. Although Arabidopsis and Camelina were harvested at species-appropriate developmental stages to prevent rhizotron overgrowth, we cannot exclude that ontogenetic differences contributed to interspecific microbiome variation. Nonetheless, the conserved enrichment of LC aliphatic GSLs at the root tip in both species suggests that axial allocation patterns are developmentally robust. Previous research has shown that GSL production and accumulation vary throughout the plant life cycle [27, 62]. Similar trends have been reported in distantly related species; for example, Huang and coworkers [52] observed a divergent pattern in microbial communities in rice and wheat grown in the same soil. However, our comparison of two closely related Brassicaceae species provides novel insight into the fine-scale and lineage-specific nature of plant–microbe interactions. Notably, the observed microbiome shifts occur in mutants defective in allocation rather than biosynthesis, reinforcing the idea that metabolite transport can function as an ecological regulatory layer independent of total metabolic output.

Together, we concluse that transporter-mediated metabolite allocation likely partakes in fine-scale chemical landscapes that likely constrain microbial colonization along the root axis. By modulating where defense metabolites accumulate, plants may flexibly tune microbial filtering across developmental zones. This perspective adds a layer to current models of plant–microbiome assembly and illustrates the importance of considering root zonation when linking specialized metabolism to microbial ecology.

## Material and Methods

### Plant material

For *Arabidopsis thaliana*, the Columbia-0 ecotype was used as the wild type reference. The *gtr1gtr2* double mutant was previously described [35]. For *Camelina sativa*, cultivar 139 (available from the Genebank of IPK Gatersleben) was used as the wild type. CRISPR-derived mutant lines G1 and M1, developed in the same genetic background, were used as described in [39]. Seeds were surface-sterilized by immersion in 70% ethanol containing 0.1% Triton X-100 for 5 minutes, followed by a 1-minute wash in 100% ethanol, and then allowed to air-dry under sterile conditions. Sterilized seeds were pre-germinated on 1/2 Murashige and Skoog (1/2 MS) medium (2.22 g□L□¹ MS basal salts, Duchefa, including vitamins and MES buffer; pH adjusted to 5.7 with KOH) after stratification for 2–3 days at 4 °C in the dark. Plants were grown under standard growth conditions: 22 °C 10h light/14h dark, 60 % humidity, 180–230 μmol m□² s□¹.

### CD-Cover setup

The CD-cover rhizotron setup was prepared as previously described [40]. To prevent root growth outside the rhizotron, the sides of each CD-cover were sealed with 3M Micropore tape. Approximately 50 g of Cologne Agricultural Soil (CAS), batch 17 from spring 2023, was used to fill each cover. Seeds were pre-germinated on 1/2 MS medium for 7 (*Arabidopsis*) and 3 days (*Camelina*) before transplantation. Two *Arabidopsis* plants or one *Camelina* plant were grown per CD-cover. The rhizotrons were positioned vertically at a 90° angle on 3D-printed racks placed inside boxes and were watered every few days until harvest. Plant material was collected after 3 weeks for *Arabidopsis* and after 7 days for *Camelina* (excluding MS germination time), to prevent root overgrowth beyond the rhizotron boundaries. This system allows for the opening of the CD-cover and complete recovery of individual root systems.

### GSL extraction and measurement

*Arabidopsis* and *Camelina* plants were grown on CD-cover rhizotrons as described above. Plants were gently removed from the rhizotrons, and roots were thoroughly washed from the soil using sterile water. Clean roots were either collected in their entirety as whole-root samples or, following the removal of lateral roots, the primary root was divided into three equal segments for section samples. Each individual root sample was measured, placed into a 0.2 mL microcentrifuge tube, flash-frozen in liquid nitrogen, and stored at –80 °C until further processing. For GSL extraction as desulfo-GSLs, the protocol described in reference [63] was followed (basic protocol 1). Samples were first homogenized using a Retsch tissueLyzer (twice for 1 minute at 30 Hz), then extracted with 85% methanol containing 10 μM of p-hydroxybenzyl GSL (pOHB, PhytoLab, cat. No. 89793) as an internal standard. Following extraction, the samples were transferred to a filter plate and washed sequentially with 70% methanol and water. Desulfation was carried out by overnight incubation with sulfatase. After elution, the resulting desulfo-GSLs were subjected to downstream analysis on an Advance UHPLC system (Bruker, Bremen, Germany) equipped with a Kinetex® XB-C18 column (100 x 2.1 mm, 1.7 µm, 100 Å, Phenomenex, USA) coupled to an EVOQ Elite TripleQuad mass spectrometer (Bruker, Bremen, Germany) equipped with an electrospray ionization source (ESI). The elution profile and mass spectrometer settings were as previously described [63] (alternate protocol 2). Glucosinolates were quantified relative to pOHB using experimentally determined response factors in a representative plant matrix.

### Bacterial and DNA isolation and amplicon sequencing

Bacterial profiling was conducted on plants grown in CD-cover rhizotrons. The specific time points for microbiome sampling are indicated in the corresponding results section. For each condition and genotype, eight individual plants grown in CAS soil were used. Large soil clumps attached to the roots were manually removed. To collect the rhizosphere fraction, cleaned roots were collected in 2 mL tubes containing sterile Milli-Q water and gently inverted several times. The resulting suspension was centrifuged, and the pellet was transferred to lysing matrix tubes for DNA extraction. For root-associated microbiota, roots underwent an additional wash to remove residual soil, then were transferred to lysing matrix tubes with screw caps (MP Biomedicals FastDNA™ SPIN Kit for Soil). For root sectioning, roots were placed on a sterile surface, measured, and divided into three equal segments. Rhizosphere and root sample processing for the sections followed the same cleaning and extraction procedures as full roots. DNA was extracted using the soil microbiome extraction protocol provided with the FastDNA™ SPIN Kit. DNA concentration was quantified using the Quant-iT™ PicoGreen™ dsDNA Assay Kit. Samples with DNA concentrations greater than 1 ng/µl were subsequently diluted to a final concentration of 1 ng/µl. Library preparation for amplicon sequencing of a c. 400-bp fragment of the V5-V7 hypervariable region of the bacterial 16S rRNA gene was carried out using the forward primer 799F 5’-AACMGGATTAGATACCCKG-3’ and the reverse primer 1193R 5’-ACGTCATCCCCACCTTCC-3’ [64, 65]. A two-step PCR protocol was used, for amplification and double-indexing with sample-specific barcodes (Supp. Table 3). Samples from *Camelina* and *Arabidopsis* were pooled separately and gel-purified for subsequent sequencing on an Illumina NovaSeq platform.

### Microbiome sequence processing and analysis

For sequence processing, a QIIME 2 pipeline was used [66]. Forward and reverse reads were first paired and demultiplexed based on barcode combinations, followed by primer sequence removal using Cutadapt [67]. The sequences were quality-checked, denoised, filtered, and amplicon sequence variants (ASVs) were generated using DADA2 [68]. Chimera detection was performed with UCHIME implemented in VSEARCH, and chimeric sequences were removed [69, 70]. Taxonomic annotation was conducted using a scikit-learn–based Naive Bayes classifier trained on the SILVA database [71]. Chloroplast and mitochondrial sequences were discarded.

### Microbiome sequence analysis

All data processing and analyses were conducted in R, with the packages magrittr and tidyverse used for data manipulation, and data visualization performed using the packages ggplot2, cowplot and gridExtra [72–76]. To minimize the influence of sequencing artifacts, amplicon sequence variants (ASVs) represented by <100 reads were removed. Rarefaction curves were generated for all samples using the vegan package to confirm sufficient sequencing depth [77]. Depending on the specific analysis, datasets were subsetted based on compartment, and subsequently rarefied to either the lowest read count per sample or, when this was below 10,000 reads, to a fixed threshold of 10,000. For datasets containing samples below this threshold, relative abundances were additionally calculated and used in downstream analyses.

Alpha diversity metrics, including ASV richness, Shannon diversity, and Pielou’s evenness were calculated for full root samples to assess the effects of species, genotype, and compartment [78, 79]. Data were tested for normality (Shapiro-Wilk test) and homogeneity of variance (Levene’s test) prior to statistical testing [80, 81]. Analysis of variance (ANOVA) followed by Tukey’s Honestly Significant Difference (HSD) test was conducted using the agricolae package [82–84].

Beta diversity was assessed both for the full root dataset and for datasets from root sectioning. Bray–Curtis dissimilarities were computed and principal coordinates analysis (PCoA) performed using the function vegan::vegdist and base R::cmdscale [85, 86]. Permutational multivariate analysis of variance (PERMANOVA) [87] was used to quantify the variance explained by experimental variables, and pairwise PERMANOVA was employed to further explore genotype and section-specific effects. In addition, datasets were aggregated at the phylum level, and after adding a pseudocount of 1, relative abundances were calculated and centered log-ratio (CLR)-transformed to address compositional data constraints. Group differences were then assessed using ANOVA followed by Tukey’s HSD test, conducted separately for each phylum.

## Supplementary figures

**Supp. Figure 1.**
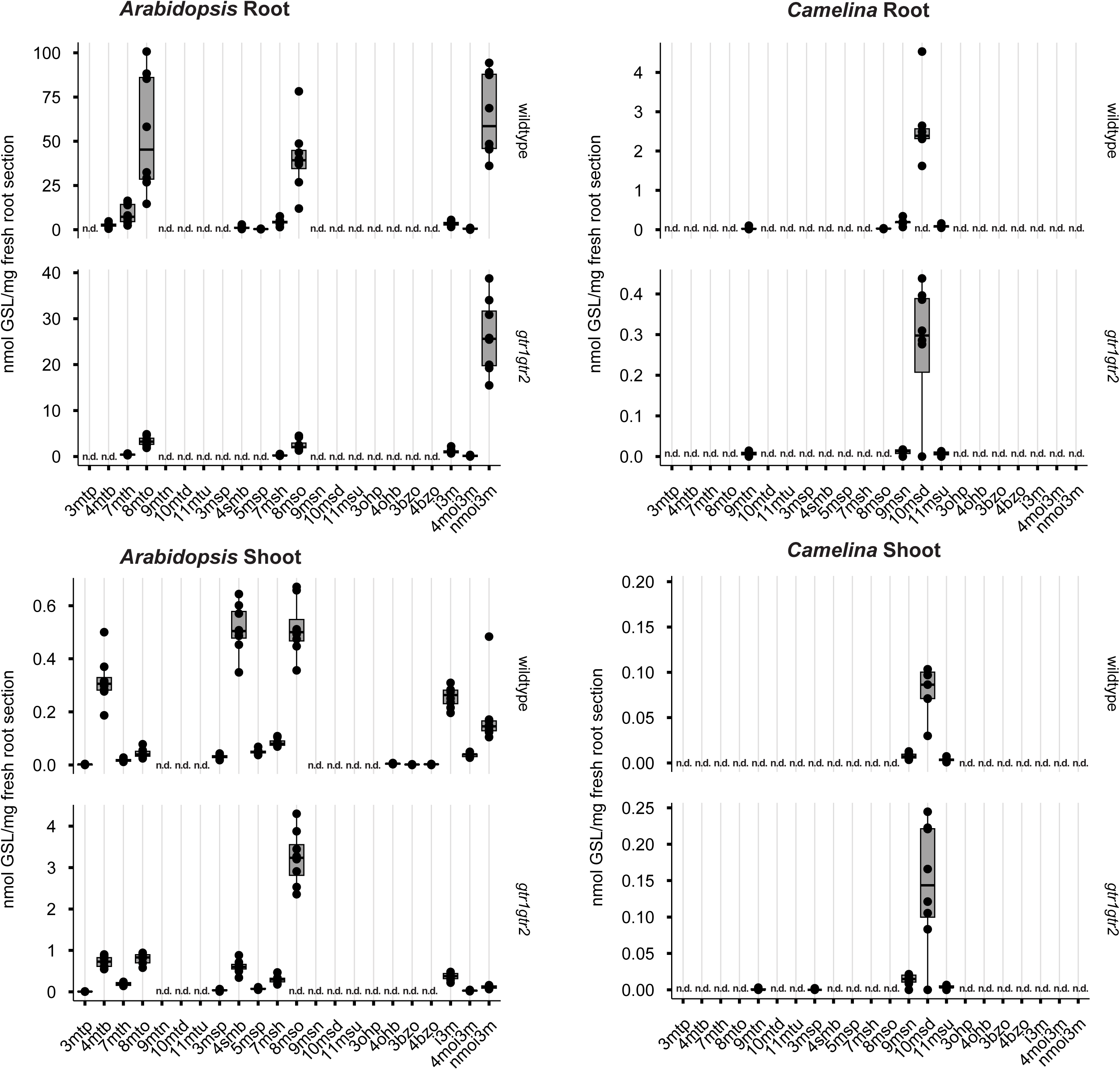
Content of individual glucosinolate types. Concentrations of individual short-chain and long-chain aliphatic and indolic glucosinolates in roots and shoots of wild type (WT) and *gtr1gtr2* mutant plants of *Arabidopsis thaliana* (*Arabidopsis*) and *Camelina sativa* (*Camelina*) grown in Cologne agricultural soil. For boxplots, the center line indicates median, dots represent data points, the box limits represent the upper and lower quartiles, and whiskers indicate maximum and minimum values. *** depicts P<0.01 in a Students t-test, n=8; n.d., not detected.

**Supp. Figure 2.**
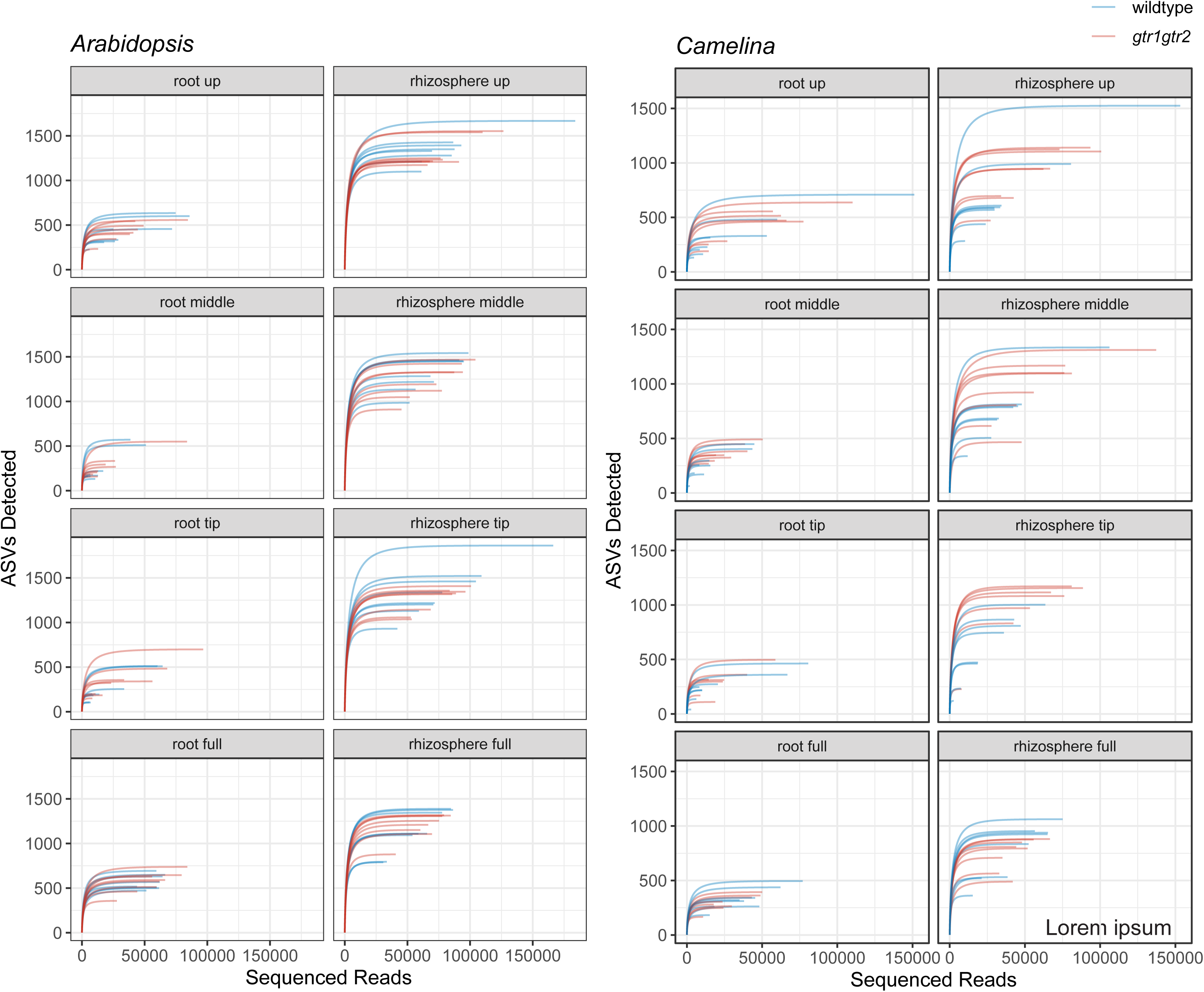
Sequencing depth of bacterial communities. Rarefaction curves showing sequencing coverage of entire roots and individual root sections of *Arabidopsis* and *Camelina* in root and rhizosphere samples.

**Supp. Figure 3.**
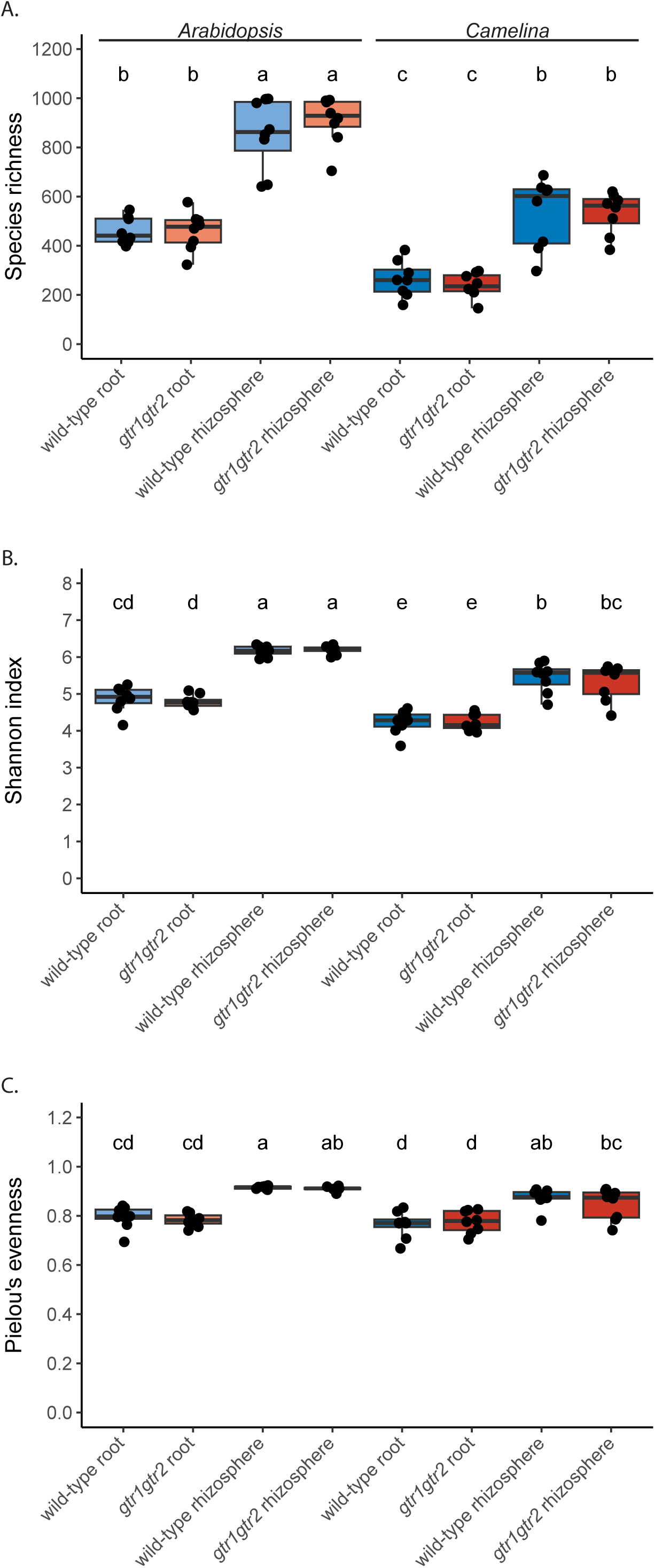
Alpha-diversity of root- and rhizosphere-associated bacterial communities. Boxplots showing (A) ASV richness, (B) Shannon diversity, and (C) Pielou’s evenness of entire roots of plants grown in Cologne agricultural soil. Statistical analysis was performed using one-way ANOVA followed by Tukey’s post hoc test Significant differences (P<0.05) are indicated by letters above the boxplots.

**Supp. Figure 4.**
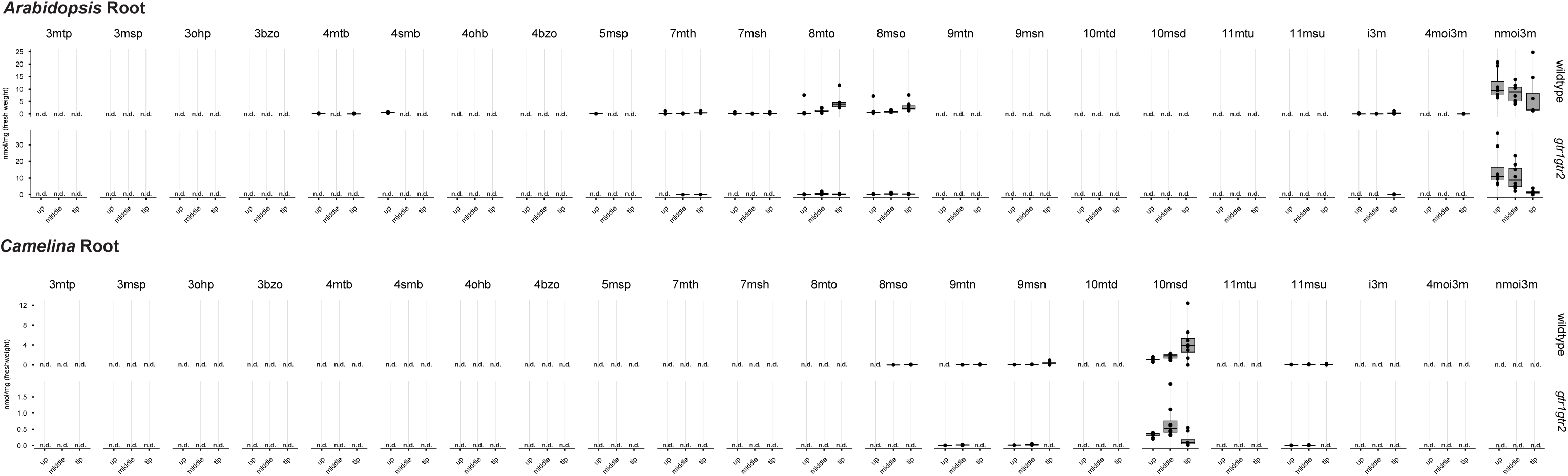
Content of individual glucosinolate types in root sections. Concentrations of individual short-chain and long-chain aliphatic and indolic glucosinolates in upper, middle and tip part of roots of wild type (WT) and *gtr1gtr2* mutants of *Arabidopsis* and *Camelina* grown in Cologne agricultural soil. For boxplots, the center line indicates median, dots represent data points, the box limits represent the upper and lower quartiles, and whiskers indicate maximum and minimum values. *** depicts P<0.01 in a Students t-test, n=8; n.d., not detected.

**Supp. Figure 5.**
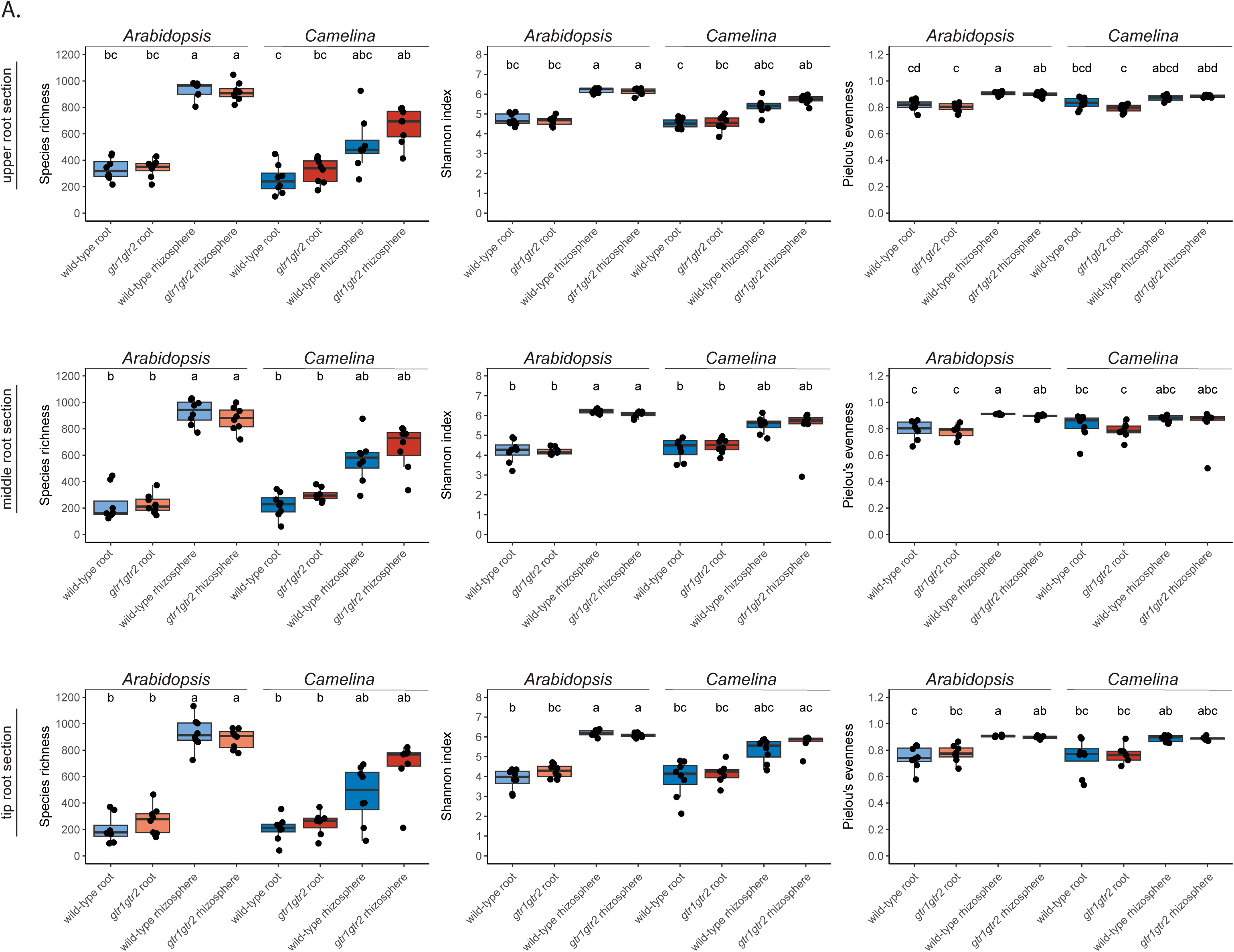
Alpha-diversity of root- and rhizosphere-associated bacterial communities across root section. Boxplots showing ASV richness, Shannon Index, and Pielou’s evenness of individual root sections of plants grown in Cologne agricultural soil. Statistical analysis was performed using one-way ANOVA followed by Tukey’s post hoc test. Significant differences are indicated by letters above the boxplots.

**Supp. Figure 6.**
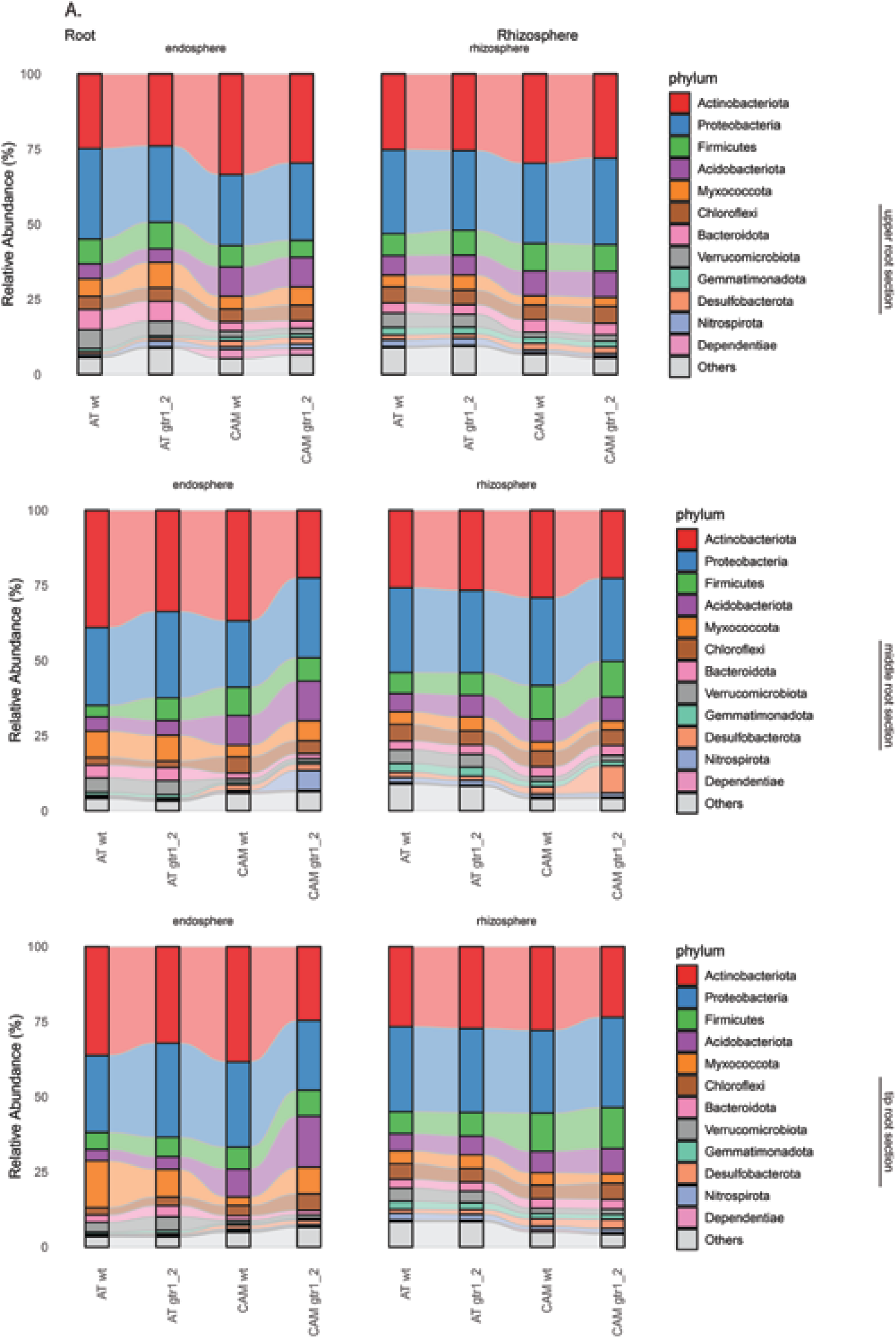
Relative abundance of most abundant bacterial phyla in roots of *Arabidopsis* and *Camelina* across rhizo-compartment, root sections, and genotype. Alluvial plots showing relative abundance of the 12 most abundant bacterial phyla across plant species and genotypes in root compartments and the three individual root sections.

## Supplementary Tables

**Supp. Table 1 | Statistical data for PERMANOVA analysis**

**Supp. Table 2 | Statistical data for ANOVA analysis**

**Supp. Table 3 | Metadata and barcodes used in this study**

## Author contributions

Conceptualization: T.G.A, L.R., A.O.R. Funding acquisition: T.G.A, A.O.R. Investigation: L.R., A.O.R. Methodology: M.B, L.R., A.O.R. Project administration: T.G.A. Supervision: T.G.A.; Visualization: A.O.R, L.R. Writing & editing: all authors.

## Acknowledgements

We wish to thank Aristeidis Stamatakis and the greenhouse team at MPIPZ for help with plant growth, Anika Kira Schroeder and Bart Boesten for technical help, and Christoph Crocoll for help with metabolic analysis. TGA also thanks the Alexander von Humboldt Foundation for funding via the Sofja Kovalevskaya program and the Max Planck Society for core funding. A.O.R was funded via a SNF postdoc mobility grant (P500PB_211108) and an EMBO Long-term fellowship (ALTF-1_2022).

## Data availability

All data are available in the main text or the supplementary materials. 16S rRNA gene amplicon reads generated in this study have been deposited at the National Center for Biotechnology Information under BioProject ID: PRJNA1426983

